# Electrical coupling between chromatophore muscle fibers allows for versatile control of chromatophore expansion in squid

**DOI:** 10.1101/2020.02.17.951715

**Authors:** Yen-Chyi Liu, Wei-Chun Wang, Bret Grasse

**Affiliations:** Neurobiology & Anatomy Department, University of Utah, Salt Lake City, UT 84112, USA; Marine Biological Laboratory, Woods Hole, MA 02543, USA; Monterey Bay Aquarium, 886 Cannery Row, Monterey, CA 93940, USA

## Abstract

Camouflage color change in cephalopods is generated by the expansion and relaxation of hundreds to thousands of chromatophore pigment organs in the skin. Individual chromatophores display color when the central pigment sac is expanded to several times its original size by a ring of 20-30 radial muscles, which are electrically coupled and are innervated by multiple motoneurons. However, mechanisms for their neuromuscular control are unclear. Here we characterize chromatophore expansion kinematics and perform simultaneous whole-cell recordings on pairs of muscle fibers of a chromatophore in squid hatchlings. We show that activity is highly correlated between muscle fibers of a chromatophore due to a high ratio of electrical coupling between all muscles for slow currents. However, fast currents are filtered and decrease rapidly further away from the muscle receiving the inputs. This low-pass filtering property of electrical coupling allows fast inputs to activate one muscle while persistent inputs spread throughout the chromatophore to synchronously activate all muscles, providing a versatile mode of control for chromatophore expansion.

## Introduction

The remarkable ability of cephalopods to rapidly alter skin coloration and patterning lies in arrays of pigment organs within the skin called chromatophores. Chromatophores consists of several cell types including a central pigment cell containing pigment granules attached to a ring of 20-30 radial muscles in a hub-and-spoke like manner [1, 2], which are directly innervated by motoneurons originating from the brain via the pallial nerve [2–4]. When chromatophore muscles contract, they significantly stretch the central pigment cell to reveal the color imparted by pigment granules. The precise control of two to four types of chromatophores of different colors (depending on species) [1, 5–7] allows the generation of a variety of patterns that can be used for camouflage or inter- and intraspecific communication [5, 8, 9].

While camouflage behavior [8–10], chromatophore ultrastructure and innervation, organization of chromatophore brain regions [4, 11–13], and chromatophore motor units [4, 14–16] have been studied, much less is known about neuromuscular control at the individual chromatophore level. Previous works have shown electrical and dye-coupling between adjacent chromatophore muscles via gap junctions and that stimulating chromatophore nerves resulted in excitatory post-synaptic potentials (EPSPs) but not action potentials [17–20]. These works shed light on the basic properties of chromatophore muscles, but mechanisms for control are still unclear: how is chromatophore expansion generated and what is the extent and function of electrical coupling?

Here we systematically examine mechanisms of individual chromatophore control with a novel dual muscle whole-cell patch clamp recording preparation in the hatchling squid *Sepioteuthis lessoniana*. First, we characterized kinematic properties of chromatophore expansions in freely-behaving animals. Next, we showed that the activity of muscle fibers of a chromatophore is extremely correlated via a high degree of electrical coupling between fibers. We then characterized the electrical coupling characteristics between muscle fibers and identified the time and distance-dependent filtering properties between fibers. Lastly, we clarified the function of electrical coupling by decoupling individual muscle fibers to examine isolated muscle responses to neural inputs. Our data suggest that fast inputs to a single chromatophore muscles can result in unidirectional expansion while sustained inputs result in omnidirectional expansion of the central pigment cell, providing a versatile mode of control for chromatophore expansion.

## Results

### Hatchling squid chromatophores make smooth and circular expansions controlled by muscles receiving highly correlated inputs

We first assessed basic expansion kinematics of chromatophores through high-speed video in freely-behaving hatchling squids (Fig. 1A). Hatchling *Sepioteuthis lessoniana* have yellow and brown chromatophores, and we chose to investigate the larger, brown chromatophores in our preparations. Multiple chromatophores on an unrestrained animal were tracked and area change, speed of area change, and circularity was calculated for each expansion/retraction event (Fig. 1B-D) (N[animals]=3, n[trials]=39). Chromatophores expanded up to twice the original area and expanded faster than retracted while maintaining a circular shape throughout the event, as was evident in calculations of circularity and aspect ratio (Fig. 1E-H). We did not observe chromatophores expanding in oblong or irregular shapes during these events, suggesting that in free swimming hatchling squids, all muscles of a chromatophore contracted synchronously throughout chromatophore expansion.

**Fig 1.**
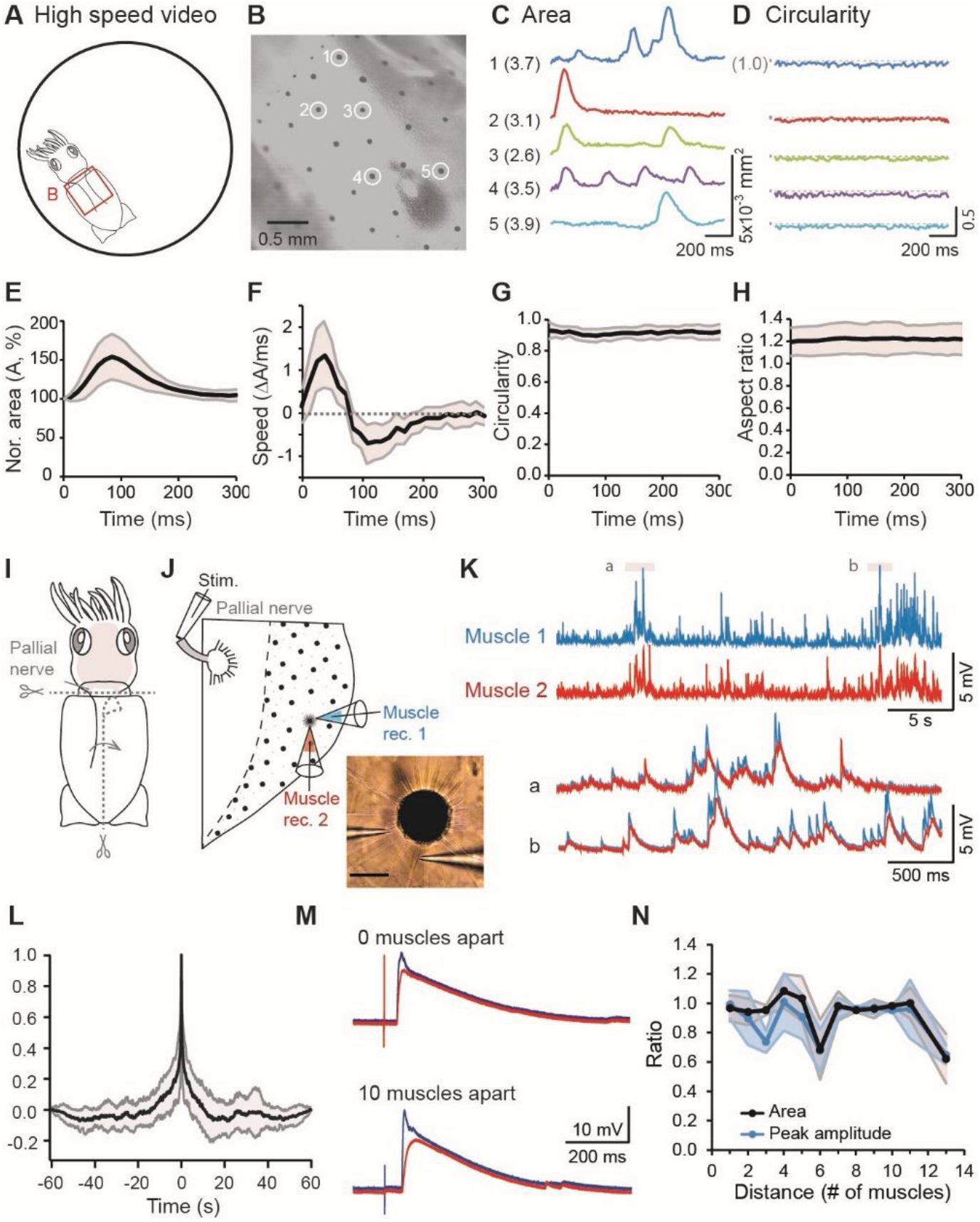
Hatchling squid chromatophores make smooth and circular expansions controlled by muscles receiving highly correlated inputs. (**A-B**) Large brown chromatophores on the dorsal mantle of a freely-behaving squid were imaged with a high-speed camera and analyzed. (**C**) The area change of five example chromatophores in B. Numbers in parentheses indicate the initial areas (unit: 10^−3^ mm^2^). (**D**) The circularity change of the five example chromatophores in B. The black dot and dashed lines indicate the location of 1, which represents a perfect circle, for each trace. (**E**) The chromatophore area change, temporally aligned to the peak of each trace and normalized to the baseline value. (**F**) The change in speed of the normalized chromatophore area in E. (**G-H**) Chromatophore circularity and aspect ratio remain constant and close to 1 during chromatophore expansions. (**I-J**) Schematic representation of preparation for dual chromatophore muscle recordings. Gray shading marks the brain and pallial nerves. The inset shows a photo of the chromatophore with two recording electrodes. Scale bar: 50 μm. (**K**) Spontaneous inputs to recorded muscle pairs are synchronous and have similar waveforms. (**L**) The cross correlation (CC) peak is close to 0s for 60s recordings of spontaneous activities. (**M**) Voltage responses of two muscles separated by different distances to pallial nerve stimulation. (**N**) Voltage responses between muscle pairs are comparable in area under curve (black) and peak amplitude (blue). The bold lines represent the means and the shading represents standard deviations for all panels.

To directly examine chromatophore control in individual muscle fibers, we performed whole-cell current-clamp recordings from two muscle fibers of a chromatophore that were separated by different numbers of muscle fibers (Fig. 1I-J). Spontaneous EPSP activity from muscle pairs of the same chromatophore were strikingly similar in overall waveform and timing, regardless the distance (number of muscle fibers) separating them (Fig.1K). Cross-correlation analyses of 60-second spontaneous recordings showed an average peak around 0 seconds (Fig. 1L) (N=13, n=13), demonstrating highly synchronized membrane activities in muscle fibers of the same chromatophore.

We next stimulated the pallial nerve with brief electric pulses via a suction electrode to investigate evoked voltage responses of muscle pairs (Fig. 1J) (N=15, n=382). A range of stimuli were applied (0.1-2 ms, 20-420 μA), but were restricted so that muscles did not contract violently to detach from the recording electrode. Muscles responded to pallial nerve stimulation with voltage depolarization that varied in amplitude with an average of 15.65 ± 8.11 mV (mean ± standard deviation) (Fig. S1A), and an action potential was not observed for our range of stimuli. Voltage waveforms of evoked responses in the recorded muscle fibers overlapped significantly with each other when superimposed, especially when muscle pairs were close together (Fig. 1M, upper panel). Evoked responses from muscle pairs further apart sometimes showed more obvious differences in peak amplitude (Fig. 1M, lower panel), however average evoked peak amplitude and area under curve across preparations were comparable between muscle pairs (Fig. 1N and S1B). Evoked responses highly correlated in timing, amplitude, and area under curve between muscles of the same chromatophore suggests synchronized control of muscle fibers within a chromatophore. Synchronized muscle control provides a mechanism for generating circular expansion of chromatophores observed in freely-behaving animals. Simultaneous high-speed video recordings during pallial nerve stimulation occasionally revealed single muscle contractions that slightly expanded the chromatophore in one direction (Fig. S1C), similar to what has been previously reported [17].

### Electrical coupling of chromatophore muscles decreases with distance and increases with the duration of input currents

How are highly-correlated voltage responses generated in different muscles within a chromatophore? We hypothesized that the electrical coupling between muscles of a chromatophore enabled inputs from a single motoneuron innervating only a few muscle fibers to generate highly-correlated responses in the rest of the muscles. To test this hypothesis, we investigated characteristics of electrical coupling between muscles of a chromatophore through dye coupling and square wave current injections of varying amplitudes. Low molecular weight fluorescent dye (Alexa Fluor 488 or 568 Hydrazide) loaded into a patch electrode was allowed to diffuse freely from a patched muscle fiber. The dye was observed to spread from the recorded muscle to adjacent muscles on either side with no directionality of spread within 15 minutes of filling (Fig. S2A). Most muscles within a chromatophore could be filled in 30-60 minutes (Fig. 2A), with exceptions where the electrode detached prematurely, or where muscles were damaged and dye did not diffuse across the damaged fibers. Fluorescent dye could be traced to the distal ends of chromatophore muscles where they sometimes branched, but was never observed to spread to another chromatophore (Fig. S2B-C).

**Fig 2.**
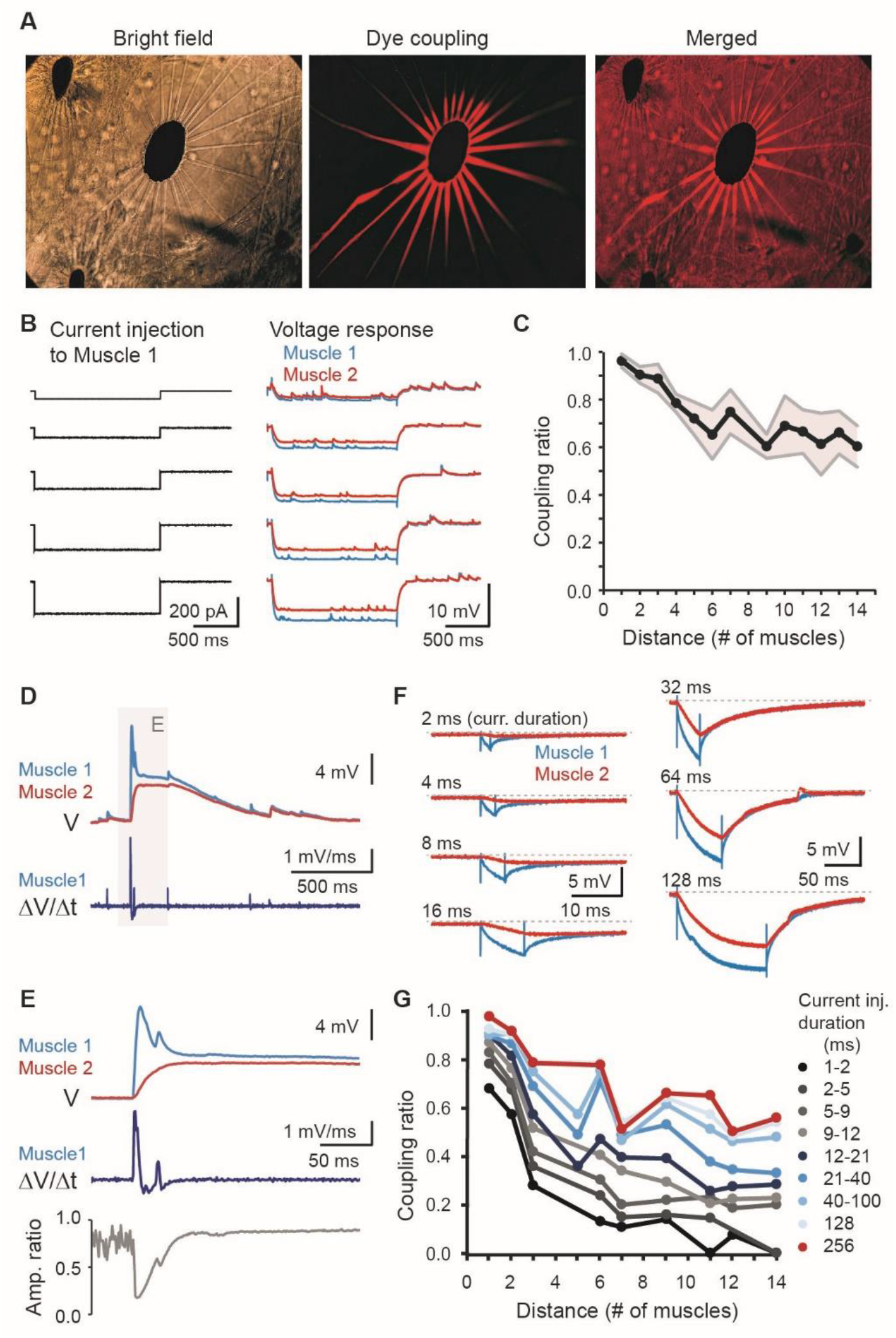
Electrical coupling of chromatophore muscles decreases with distance and increases with the duration of input currents. (**A**) Alexa Fluor 568 was introduced into a single chromatophore muscle fiber through a patch pipette and spread to all muscle fibers within 45 mins in this trial. (**B**) Example voltage responses of muscle 1 (blue) and 2 (red) to 1.5 second hyperpolarizing current injections with varying amplitudes to muscle 1. (**C**) The coupling ratio (muscle 2/ muscle 1) decreases with distance between chromatophore muscles. The bold line represents the mean and the shading represents the standard deviation. (**D**) Example traces of voltage responses in muscle pairs 10 muscles apart (note the lack of a fast component in muscle 2 compared to muscle 1). The time derivative of the voltage trace of muscle 1 is shown in the bottom panel. (**E**) An expanded view of the shaded area in D. The amplitude ratio (muscle 2/ muscle 1) decreases when the rate of voltage change in muscle 1 increases. (**F**) Example voltage responses of muscle 1 and 2 to injecting 400 pA hyperpolarizing currents with different durations to muscle 1. (**G**) In response to current injections of the same duration, the coupling ratio decreases with muscle distance as in C. Shorter duration currents are generally less well-coupled than longer duration currents, especially at further distances apart (black line).

We next examined the degree of electrical coupling between muscle fibers at varying distances through injecting 1.5 second hyperpolarizing square wave currents into one muscle and calculating the ratio of the voltage responses at steady state between the injected muscle and the second muscle (Fig. 2B) (N=20, n=209). This “coupling ratio” was extremely high (> 90%) for muscles close to one another, and began to decrease gradually after three muscles (Fig. 2C). However, even when the two muscle fibers were furthest apart on opposite sides of the chromatophore, the coupling ratio at steady state remained high (> 60%) (Fig. 2C).

Even though a high level of electrical coupling was observed, stimulating the pallial nerve sometimes resulted in evoked responses in one muscle that was slightly larger in amplitude and had a fast component compared to the other (Fig. 1M), suggesting differential filtering of input to the two muscle fibers. To investigate the filtering properties of muscle fibers, we took the first derivative of the voltage waveform with a fast component and compared the rate of voltage change to the amplitude ratio between the muscle fibers (Fig. 2D-E). We found that higher rates of voltage change corresponded to lower amplitude ratios, suggesting that the input current had been low-pass filtered in the muscle with the smaller response (Fig. 2E). To investigate this characteristic systematically, we conducted a series of current injections of the same amplitude that varied in duration and measured the maximum coupling ratio between muscle fibers at varying distances apart (Fig. 2F-G) (N=7, n=85). Current injections of shorter durations resulted in low electrical coupling between muscle fibers, and the ratio increased with longer duration currents (Fig. 2F-G). This was true even when two muscle fibers were adjacent to one another, and levels of electrical coupling were lower for shorter duration currents. Thus the coupling ratio for shorter duration current inputs are lower compared to long duration current inputs at the same distance and decrease faster over longer distances.

### A chromatophore muscle likely receives inputs from multiple motoneurons and through electrical coupling from other muscles in the same chromatophore

Due to the low-pass filtering property of electrical coupling between chromatophore muscle fibers, it seems unlikely that voltage responses of similar amplitudes in different muscle fibers were generated solely through electrical coupling. We tested whether electrical coupling alone was sufficient to generate comparable waveforms in two muscle fibers by simulating neuronal input to only one muscle fiber using a natural input waveform and recording the waveform response in a second muscle (Fig. 3A-D) (N=9, n=78). First, with both muscles under current-clamp, we stimulated the pallial nerve multiple times at the same setting for voltage responses (Fig. 3C). Then, one muscle (muscle 1) was switched to voltage-clamp to record the current response to the same pallial nerve stimulus (Fig. 3A). The current polarity was reversed and used as a template for current injection back into muscle 1 (Fig. 3B). The voltage responses to these template current injections were then monitored for both muscles under current-clamp (Fig. 3D). The average peak amplitude of voltage responses elicited by pallial nerve stimulation and template current injection for both muscles were analyzed and normalized to the average amplitude of muscle 1 voltage response to pallial nerve stimulation (Fig. 3E). We found that average voltage responses of muscle 1 to template current injection were comparable or often larger than the voltage response of muscle 1 to pallial nerve stimulation (Fig. 3E). This was likely due to choosing the largest evoked current as the template for current injection. As expected from previous results, the voltage response of muscle 2 to nerve stimulation was comparable to muscle 1 to nerve stimulation and only decreased at farther distances (Fig. 3E). More importantly, voltage responses of muscle 2 to template current injections in muscle 1 were significantly smaller (p<0.05, Mann-Whitney test) compared to voltage responses of muscle 2 to nerve stimulation (Fig. 3E). Likewise, the ratio of both the peak amplitude and area under curve between muscle 1 and 2 decreased significantly for template current injections compared to pallial nerve stimulation (Fig. 3F). These results demonstrate that the input to a single muscle fiber is insufficient to generate comparable voltage responses in another, regardless the distance. This is likely due to the shape of the input current waveform, causing it to be low-pass filtered through electrical coupling. This low-pass filtering property of electrical coupling between muscles provides a possible explanation for occasionally observed single muscle contractions in a chromatophore (Fig. S1C). A large input of short duration to only one muscle fiber would activate that fiber, but due to electrical filtering, would not be sufficient to activate adjacent fibers. Since low-pass electrical filtering between adjacent muscles prevents comparable voltage responses in multiple muscles to the input to a single muscle fiber, it seems likely that motoneurons innervate multiple muscles of a chromatophore and provide input to separate fibers.

**Fig 3.**
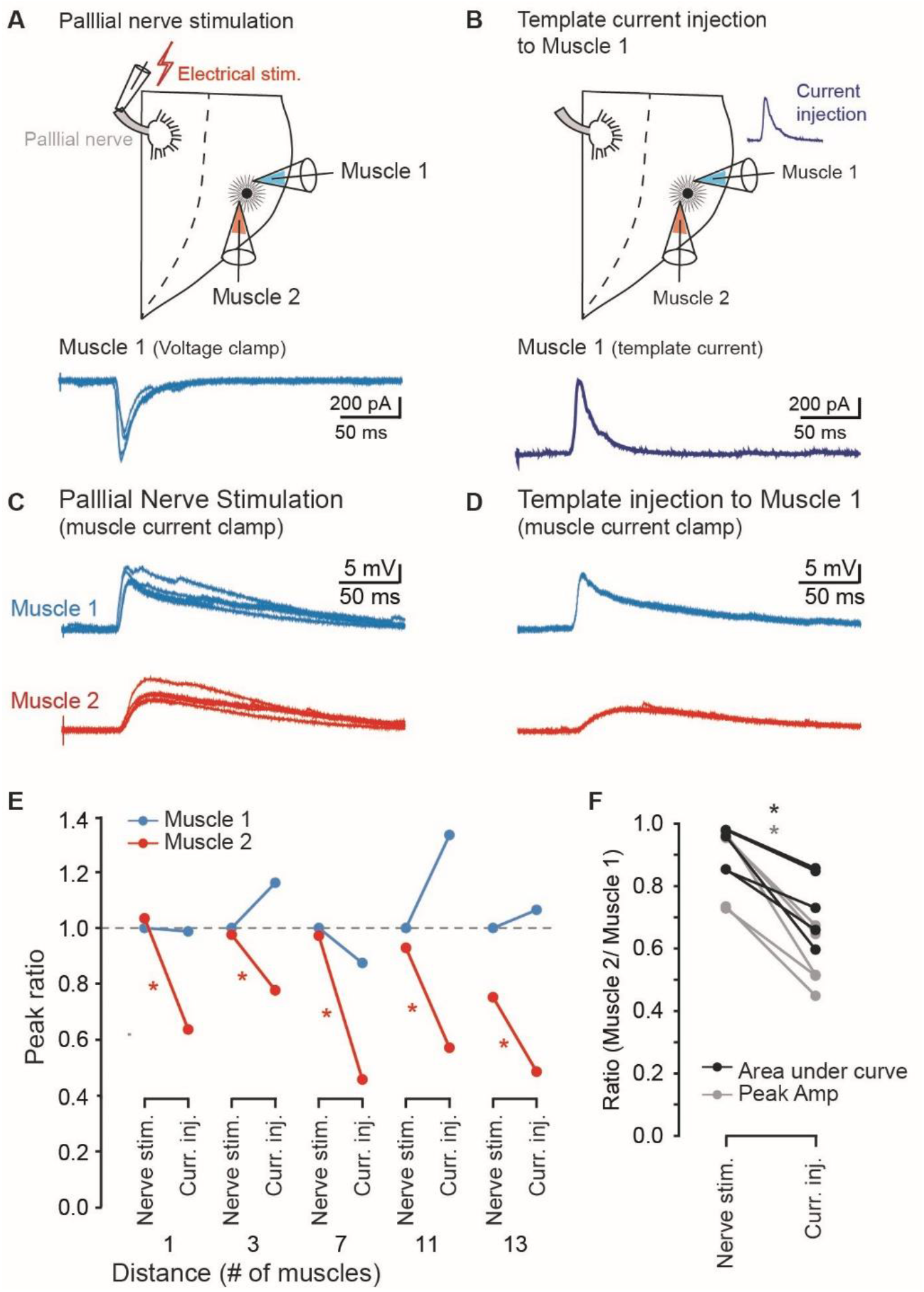
A chromatophore muscle likely receives inputs from multiple motoneurons and through electrical coupling from other muscles in the same chromatophore. (**A**) Pallial nerve stimulation elicited current inputs in the recorded chromatophore muscle (Muscle 1). (**B**) One of the current traces (usually the largest) was inverted and used as a template for current injection applied back to Muscle 1. (**C**) Example voltage responses elicited by pallial nerve stimulation in two chromatophore muscles (Muscle 1 and 2) under current clamp. (**D**) Example voltage responses of Muscle 1 and 2 in response to template current injection to Muscle 1. (**E**) The peak amplitude of voltage responses of recorded chromatophore muscles at different distances, normalized to the peak amplitude of Muscle 1 response to pallial nerve stimulation. *p<0.05, Mann-Whitney test. (**F**) Both the peak amplitude (gray) and area under curve (black) ratios (Muscle 2/ Muscle 1) elicited by pallial nerve stimulation are significantly larger than the ratios in response to template current injection to Muscle 1. Recordings from different muscle distances are pooled together (*p < 0.05, paired t test).

### The activity of isolated chromatophore muscles is not synchronous with coupled muscles

To investigate muscle activity in the absence of electrical coupling, we applied gap junction blockers octanol and 18β-glycyrrhetinic acid in attempt to disrupt electrical coupling between chromatophore muscle fibers. Neither blocker at varying concentrations could reliably decrease electrical coupling in our preparation (Fig. S3). Since dye coupling was disrupted by damaged muscle fibers in previous experiments, we attempted the same approach to disrupt electrical coupling between muscles. Muscle fibers on either side of the target muscle were physically broken with a pipette, and removed via a suction electrode (Fig. 4A). We recorded from the isolated muscle fiber and a separate muscle that remained coupled with other muscles in the chromatophore (Fig. 4B) (N=5, n=73). 1.5 second hyperpolarizing current injections to the coupled muscle did not result in a voltage response in the isolated muscle, demonstrating that electrical coupling to the isolated muscle had been disrupted (Fig. 4C). Spontaneous activity in isolated muscles did not match coupled muscles in timing, amplitude, or waveform (Fig. 4D). Average cross-correlograms of 60-second spontaneous activity traces between isolated and coupled muscles revealed no clear peaks (Fig. 4E) (N=8, n=8). Evoked responses to pallial nerve stimulation also differed drastically for isolated and coupled muscle fibers (Fig. 4F). Stimuli of the same amplitude and duration in different trials of the same animal generated comparable voltage response waveforms in the coupled muscle, but the isolated muscle sometimes responded with multiple peaks, or did not respond at all (Fig. 4F). Comparisons between coupled and isolated muscle responses to pallial nerve stimulation revealed that the average membrane time constant decreased significantly in the isolated muscles, the latency from stimulus to response peak showed a slight but significant decrease, but peak amplitude was not significantly different (p<0.05, Wilcoxon matched pairs test) (Fig. 4G-I). These data show the importance of electrical coupling between muscle fibers to ensure synchronous inputs to adjacent muscles and suggest electrical coupling as a mechanism of smoothing out current inputs from different sources.

**Fig 4.**
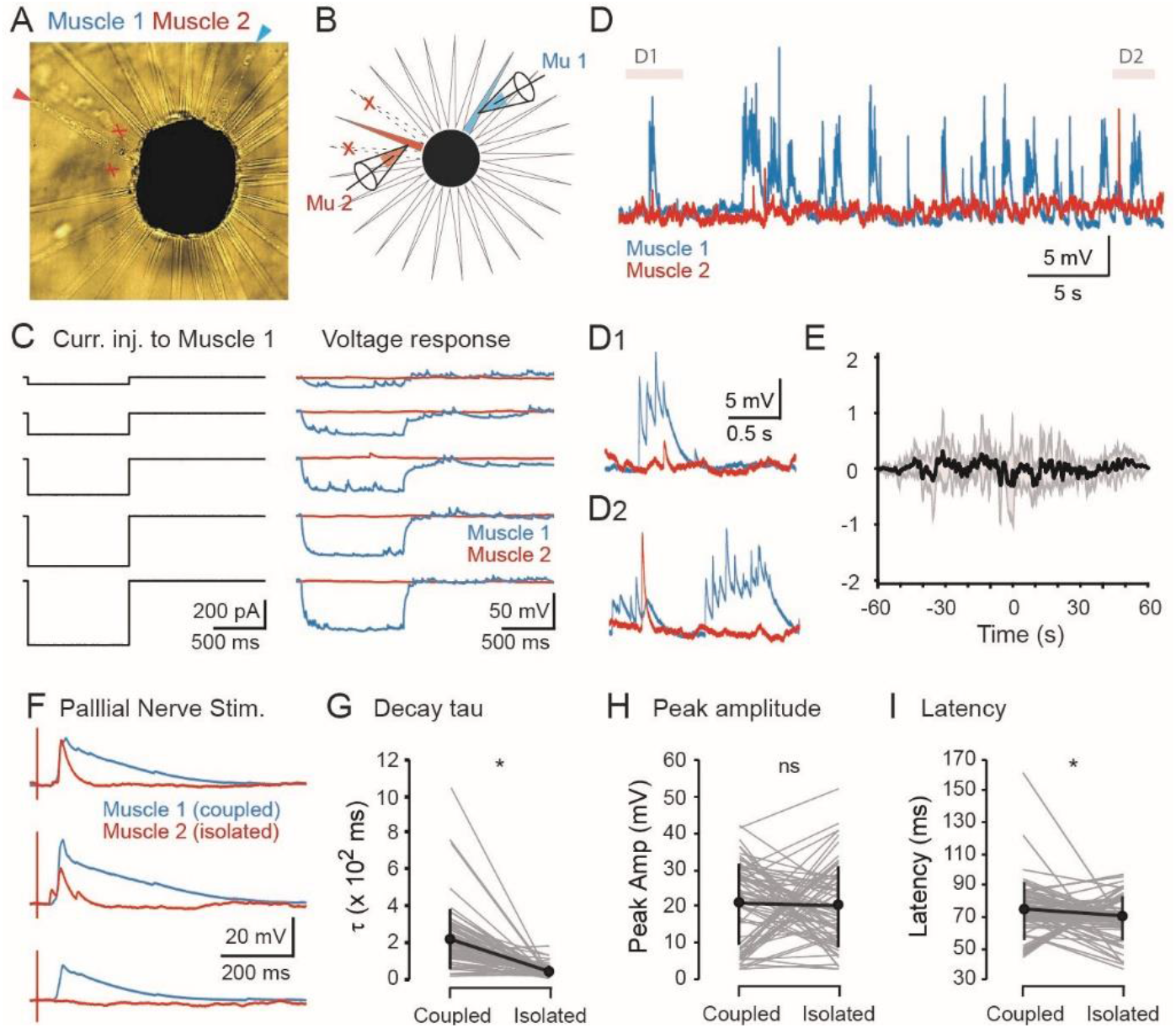
The activity of isolated chromatophore muscles is not synchronous with coupled muscles. (**A-B**) Photo and schematic of an isolated chromatophore muscle (Muscle 2, red). Muscle 1 (blue) is electrically coupled to other muscles of the chromatophore. (**C**) No electrical coupling is observed in the isolated muscle (Muscle 2) during hyperpolarizing current injections to Muscle 1. (**D**) Spontaneous activities in the isolated muscle (Muscle 2) are fewer and are not synchronous with the coupled muscle (Muscle 1). D1 and D2 show expanded views for the shaded areas in D. (**E**) There is no clear cross correlation peak for 60s spontaneous activity in simultaneous recordings of Muscles 1 and 2. The bold line represents the mean and the shading represents the standard deviation. (**F**) Example voltage responses elicited by pallial nerve stimulation of the same amplitude and duration for the same coupled (Muscle 1) and isolated (Muscle 2) muscle pair. (**G-I**) Voltage responses in the isolated muscles have shorter voltage decay time constants (tau) and shorter latencies from stimulus to peak of response. Peak amplitude was not significantly different (ns) between isolated and coupled muscles. Gray lines: individual muscle pairs; black line: mean ± standard deviation. *p<0.05, Wilcoxon matched pairs test.

## Discussion

Our results suggest a versatile mode of motor control through intrinsic properties of electrical coupling between muscles in the squid chromatophore. Fast current inputs to a single chromatophore muscle fiber may be sufficient to generate a contraction in that muscle, yet due to the input waveform being low-pass filtered, would not be sufficient to activate muscles at farther distances thus expanding the chromatophore in one direction (Fig. 5A). Alternatively, sustained inputs to one muscle fiber will allow current to spread throughout the chromatophore and cause synchronous activation of all muscles to generate circular expansion in all directions (Fig. 5B). Similarly, simultaneous current inputs to multiple chromatophore muscles would also likely generate circular expansion of the chromatophore (Fig. 5C). Unidirectional chromatophore expansion may be useful for generating patterns with high spatial resolution, while synchronous activation of muscles in a chromatophore is likely a more efficient method in the display of color and reducing the complexity of neural control. Our observations in freely-behaving, uninjured hatchling squids support omnidirectional chromatophore expansion, however our observations in semi-intact preparations (Fig. S1C), adult animals, and other cephalopod species suggest unidirectional expansion can also happen [17]. More work is needed to determine which of these methods are utilized, or if a combination of these methods is used for generation of specific patterns. Further studies on chromatophore motoneuron activity and projection patterns are also required to clarify the mechanisms of chromatophore control.

**Fig 5.**
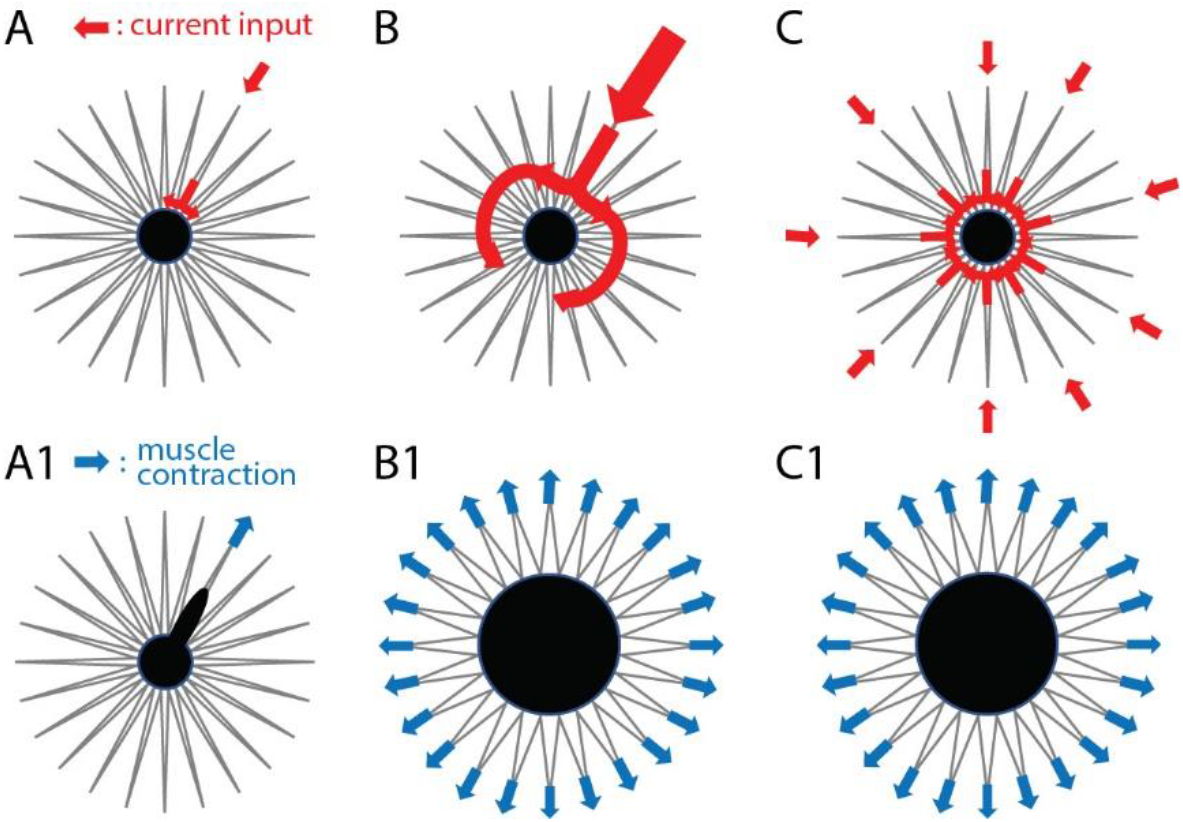
Model of a versatile control scheme for chromatophore expansion. (**A**) Due to low-pass filtering characteristics of muscles of a chromatophore, a fast current input to one chromatophore muscle fiber would be filtered out in other muscles, resulting in contraction of only the muscle receiving the input, pulling the pigment cell in one direction and causing unidirectional expansion of the chromatophore as shown in A1. (**B-C**) Omnidirectional expansion of a chromatophore may be generated by either a sustained input to one muscle (**B**) or simultaneous inputs to multiple muscles of a chromatophore (**C**). Both modes of current input result in activation of multiple muscle fibers and expansion of the pigment cell in a circular manner as shown in B1 and C1. Red arrows represent current inputs while blue arrows represent muscle fiber contraction.

## Materials and Methods

### Animals

Eggs of *Sepioteuthis lessoniana* were kept in a closed, recirculating seawater system until hatching naturally. Artificial seawater, Instant Ocean® artificial salt mixture, was utilized for this system and acceptable water quality [21] was maintained through the use of filter pad medium, activated carbon, and biofilter media. Egg capsules were suspended in the water column using zip ties and fishing line and moderate flow was directed over the eggs to promote oxygenation to the developing embryos. Non-viable eggs and spent eggs capsules were removed and discarded quickly to avoid fouling the system. Animals were removed for experimentation within 48 hours post hatching.

### High Speed Video and Analysis

A single free-swimming squid hatchling was placed in a 10 cm diameter plastic petri dish with sea water for high-speed video recordings. Spontaneous expanding and contracting of the chromatophores were imaged at 250 frames per second using a Photonics Fastcam (Photron).

The larger, brown chromatophores were chosen for analysis due to their contour being more easily identifiable than the smaller yellow chromatophores. The *Analyze Particle* function of Image J was used to detect the outline of chromatophores on every third frames, and the time series of their area, circularity, and aspect ratio was exported. Speed of area change of each chromatophore was calculated as the chromatophore area difference of consecutive frames divided by the interval between analyzed frames. Both circularity and aspect ratio were used to represent the roundness of the chromatophores. Circularity (*4π* × *[Area]/[Perimeter]*^*2*^) of one indicates a perfect circle, and aspect ratio represents the ratio of the major and minor axes of an ellipse fitted to the shape of chromatophores. For presenting the average trend of a single expansion-contraction from multiple chromatophores, the area of chromatophores was normalized to their baseline size before expansion and temporally aligned to the maximum expansion.

### Electrophysiology and Analysis

Hatchling squid were anesthetized in artificial sea water with 0.1% Tricaine and moved to a Sylgard bottom recording chamber containing artificial sea water (composition in mmol/L: 422.98 NaCl, 8.99 KCl, 9.25 CaCl_2_·2H_2_O, 22.92 MgCl_2_·6H_2_O, 25.52 MgSO_4_·7H_2_O, 2.14 NaHCO_3_; adjusted to pH 7.8 with NaOH, ~ 1200 mOsm, Marine Biological Laboratory Formulae and Methods V). The ventral mantle was cut at the midline, unfolded and stabilized with customized tungsten pins. Internal organs were removed and the dorsal mantle was cut at the midline, giving two halves, and the head and brain was severed, leaving the pallial nerve intact. The mantle musculature was carefully peeled back from the skin to preserve as much of the nerves as possible and pinned down to expose the underlying chromatophores. Sheath cells around the muscle fibers were removed with a suction electrode to access chromatophore muscle fibers for recordings (Figure 1I).

Recordings were performed under constant perfusion of chilled sea water (~12°C) [17] using a Harvard Apparatus SC-20 inline cooler with CL-100 temperature controller. Dual whole-cell current clamp recordings were performed on muscle fibers of the same chromatophore at different distances (separated by different numbers of chromatophore muscles) to examine spontaneous and/or evoked membrane potential changes to direct current injection into the recorded muscle cell or pallial nerve stimulation (Figure 1J-K). The resistance of recording pipettes was 2-5 MΩ. The recording pipette was backfilled with patch solution (composition in mmol/L: 140 KCl, 696 glucose, 1 CaCl_2_, 11 EGTA, 10 HEPES; adjusted to pH 7.4 with KOH, ~ 1200 mOsm, [22]). The pallial nerve was stimulated with a glass suction electrode and the stimulus duration and amplitude (0.1-2 ms, 20-420 μA) was adjusted to elicit depolarizations in the recorded chromatophore muscle that did not lead to violent muscle contraction and subsequent loss of recordings.

Cross-correlation (CC) was analyzed for 60 second simultaneous recordings of spontaneous activity from two chromatophore muscles. The CC peak lag (temporal interval between the CC peak and 0) was correlated to the distance of the recorded muscles. The coupling ratio of two chromatophore muscles was defined as the ratio of the maximum voltage change in each muscle cell in response to current injection to one of the two muscles under current clamp.

Whole-cell muscle recording data were acquired with a HEKA EPC10 double patch clamp amplifier controlled by HEKA Patchmaster software (Harvard Bioscience) with 100 kHz sampling rate. Data were analyzed using Igor Pro (WaveMetrics), Clampfit (Molecular Devices), and DataView [23].

### Dye Coupling

Red and green fluorescent dye (Alexa Fluor 568 and Alexa Fluor 488, final concentration 50 μmol/L) was included in the recording pipette to fill the recorded chromatophore muscle and identify surrounding muscles that were coupled to the recorded muscle.

### Statistics

All data were tested for normality to determine whether parametric or nonparametric statistic should be used. For comparing means between two groups, paired t-test was performed for parametric and Wilcoxon matched pairs test was performed for nonparametric data. Multiple comparisons were performed using one-way ANOVA, followed by post hoc Tukey-Kramer test. Correlations were examined by Spearman’s rank test. Statistical analysis was performed with Microsoft Excel and StatPlus Professional (AnalystSoft).

## Acknowledgements

We would like to thank Drs. Gail Mandel and Paul Brehm for guidance and support of this research. We would like to acknowledge the Monterey Bay Aquarium management team for supporting this collaboration.

## Supplementary Figures

**Figure S1.**
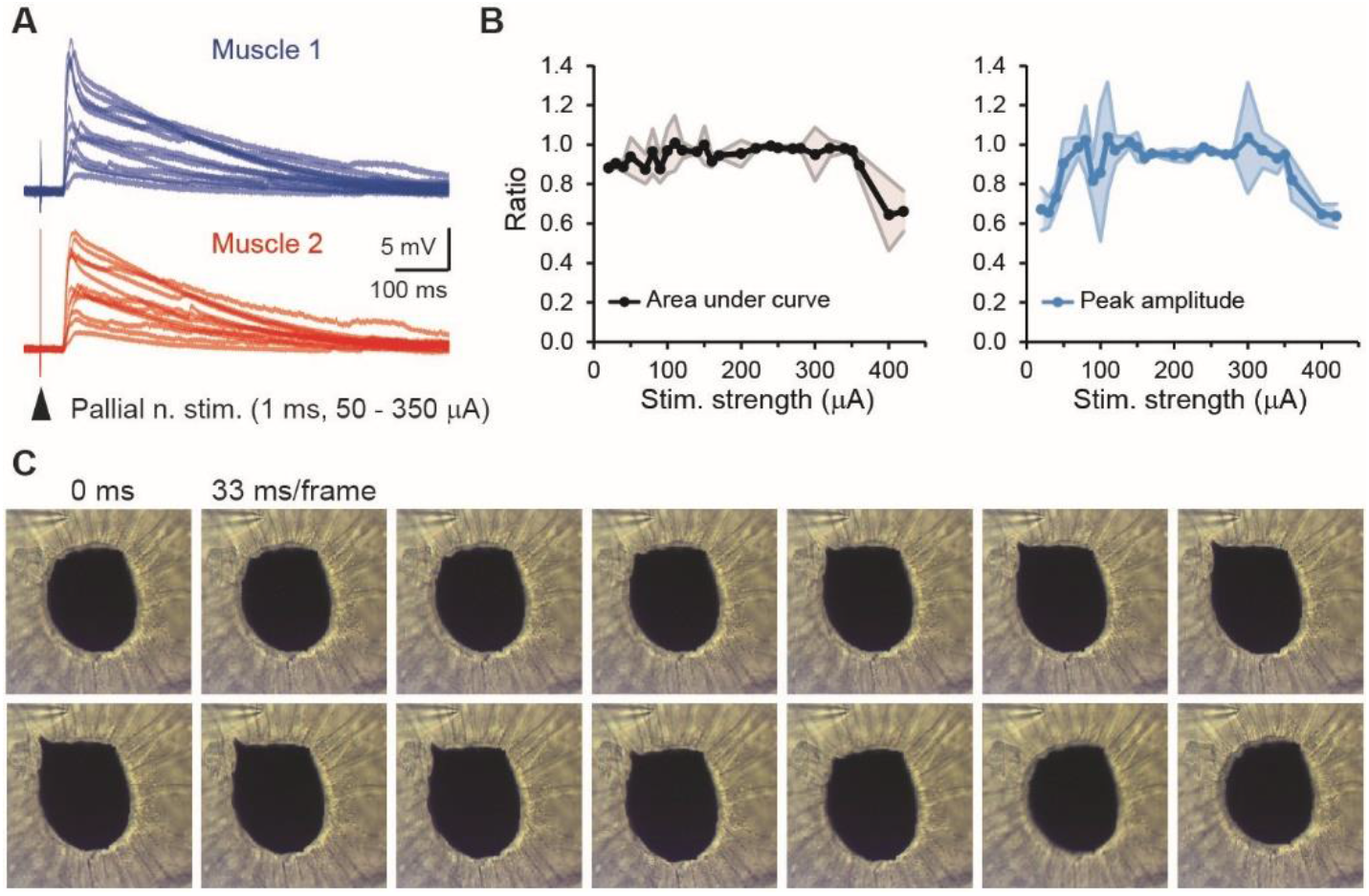
Stimulating the pallial nerve causes comparable voltage responses in chromatophore muscle cells that can sometimes generate single muscle contractions. Related to Figure 1. (**A**) Stimulating the pallial nerve at various strengths generates variable voltage responses in muscle fibers of the same chromatophore, but not action potentials. (**B**) Peak amplitude and area under curve for muscle pairs of the same chromatophore are generally comparable for varying stimulus strengths. (**C**) Single muscle contractions can sometimes be observed during pallial nerve stimulus trials where other muscles of the chromatophore do not contract, thus elongating the chromatophore in one direction.

**Figure S2.**
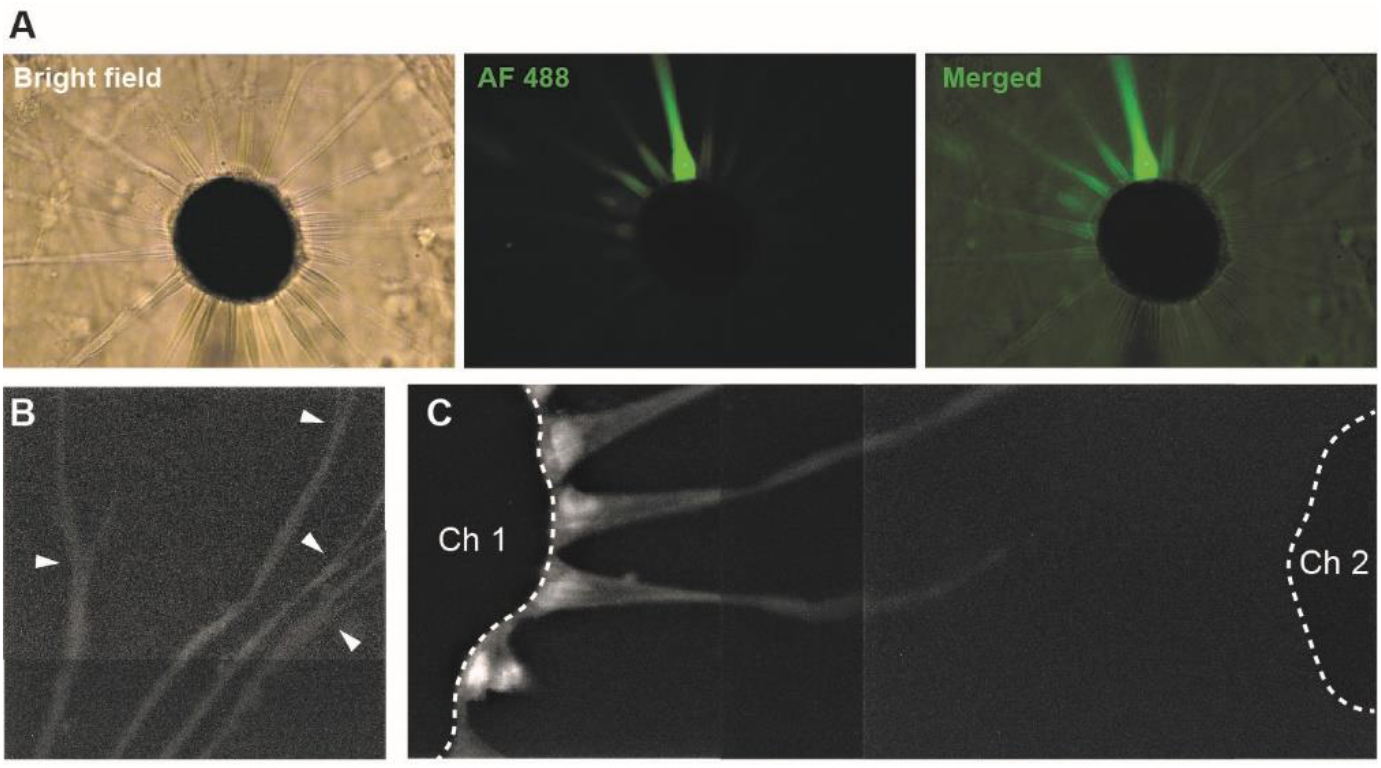
Florescent dye spreads bidirectionally and fills muscles to distal branches but do not spread to other chromatophores. Related to Figure 2. (**A**) Alexa Fluor 488 green florescent dye can be observed to spread from the original muscle (brightest muscle fiber) in both directions. We never observed dye diffusion in only one direction in our trials. Photos were taken ~15 minutes after start of fill when dye has not yet diffused to all muscles. (**B**) Following muscle fibers to distal ends often revealed branching (white arrowheads), but (**C**) florescent dye was never observed to spread from one chromatophore muscle to another. Ch1: a chromatophore where dye was allowed to fill muscles fibers; Ch2: a neighboring chromatophore with muscle fibers in close proximity to the first chromatophore, but no dye was observed to cross. Dashed white lines delineate boundaries of chromatophore pigment sac. A series of overlapping images was stacked to generate panels B and C.

**Figure S3.**
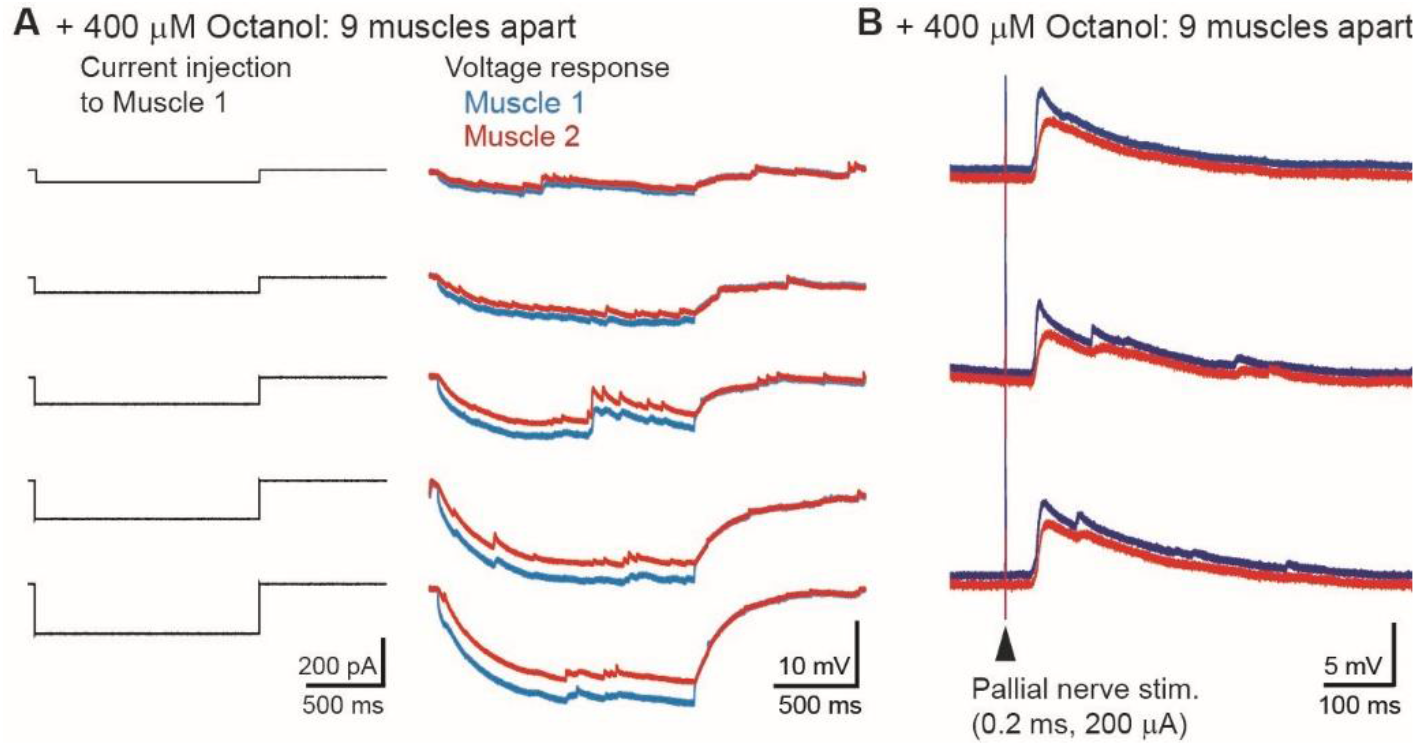
Gap junction blockers did not reliably reduce electrical coupling between chromatophore muscle fibers in our preparation. Related to Figure 4. (**A**) Addition of 400 μM octanol to the solution did not block electrical coupling between muscles that were separated by 9 other muscles for 1.5 sec hyperpolarizing current injections of varying amplitudes. (**B**) Voltage responses to pallial nerve stimulation was also comparable in amplitude for the same muscle pairs as in A.

